# Multiple focal pulvinar projection fields in the macaque cortex

**DOI:** 10.1101/2023.11.20.567885

**Authors:** Mathilda Froesel, Simon Clavagnier, Quentin Goudard, Qi Zhu, Wim Vanduffel, Suliann Ben Hamed

**Author notes:** Corresponding authors: Mathilda Froesel; Suliann Ben Hamed.

## Abstract

The pulvinar, the largest nucleus of the thalamus, is functionally heterogeneous and involved in multiple cognitive functions. It has been proposed to act as a functional hub of cortical processes due to its extensive reciprocal connectivity with the cortex. However, its role in cognition is not fully understood yet. Here, we posit that an improved understanding of its functional connectivity with the cortex is needed to better capture the cognitive functions of this nucleus. To address this question, we characterize the pulvino-cortical functional connectivity along the ventro-dorsal, antero-posterior and medio-lateral axes, using awake resting state data from ten adult macaques. We first report two global cortical functional connectivity gradients along the antero-posterior and ventro-dorsal pulvinar gradients, that match remarkably well the structural connectivity gradients described by anatomical approaches. In addition to these global gradients, multiple local cortical pulvinar projection fields can be identified at the sulci level such as in the lateral sulcus (LS), the intraparietal sulcus (IPS), the principal sulci (PS) and the anterior cingulate cortex (ACC). For most sulci, we show that functional pulvino-cortical projection fields follow the major anatomical axis of these different sulci (e.g. the ventro-dorsal axis for the LS and the antero-posterior axis for the IPS). Other sulci, such as the superior temporal sulcus, the posterior cingulate cortex or the central sulcus, display multiple projection fields from the pulvinar. Although substantial inter-individual differences exist, the general functional connectivity patterns are remarkably consistent across hemispheres and individuals. Overall, we propose that these multiple pulvinar projection fields correspond to a fundamental principle of pulvino-cortical connectivity and that a better understanding of this connectional organization will shed light on the function of pulvino-cortical interactions and the role of the pulvinar in cognition at large.

## Introduction

The pulvinar, the largest and most posterior nucleus of the thalamus, is characterized by widespread anatomical connections with the rest of the brain. Based on the highly complex cortico-pulvino-cortical connectivity and the pulvinar’s synaptic organization with typical feedback and feedforward projections, the pulvinar has been proposed to act as a functional hub or modulator of cortical processes (Benarroch, 2015; Rouiller & Welker, 2000; Saalmann & Kastner, 2011; Sherman, 2007; Sherman & Guillery, 2006; Shipp, 2003). The anatomy of this higher-order subcortical nucleus is also very heterogeneous. It is classically divided in several sub regions based on their specific cytoarchitectonic and chemoarchitectonic properties, namely the anterior, medial, lateral and inferior subdivisions of the pulvinar, each containing even smaller subdivisions (Gutierrez et al., 1995, 2000; Olszewski et al., 1952; Stepniewska & Kaas, 1997; Walker, 1938). For example, the medial pulvinar is divided in medio-lateral and medio-medial nuclei, the lateral pulvinar is divided in a ventro-lateral and dorso-lateral pulvinar. The inferior pulvinar is divided in the posterior, medial and central inferior pulvinar. In humans, functional pulvinar parcellations have mostly relied on meta-analyses of task-related neuroimaging studies and resting state-based clustering analyses (Barron et al., 2015; Guedj & Vuilleumier, 2020). Both studies describe five pulvinar functional clusters, though they do not locate them at the exact same anatomical position. The meta-analysis defines an inferior, lateral, medial, anterior and superior pulvinar cluster (Barron et al., 2015) while the resting state analysis splits the pulvinar into a dorso-medial, ventro-medial, lateral, anterior and inferior cluster (Guedj & Vuilleumier, 2020). These results demonstrate that, functionally and anatomically, the subregions of the pulvinar are complex to define.

In the present study, we analyzed functional pulvino-cortical connectivity at the spatial resolution of individual fMRI voxels along the ventro-dorsal, antero-posterior and medio-lateral axis. Based on the literature, we predict that the pulvinar follows a functional connectivity gradient with the cortex that matches the anatomical pulvino-cortical connectivity organization, namely a ventro-dorsal pulvinar gradient that maps onto an antero-posterior cortical functional gradient (Froesel et al., 2021). We also predict locally-specific topographically organized projections from the pulvinar to the cortex. Such local pulvinar projection patterns have been reported in specific cortical regions such as the superior temporal sulcus or MT (Grimaldi et al., 2016; Mundinano et al., 2019). To investigate both global and local functional connectivity patterns between the pulvinar and the cortex, we performed resting state fMRI analyses from ten awake fixating monkeys using single voxels as seeds for whole brain functional connectivity analyses. We report that, globally, as predicted by the literature, the pulvino-cortical connectivity mainly follows a ventro-dorsal and antero-posterior gradient. Most importantly, we additionally show multiple focal functional pulvinar projection fields at the cortical level, mostly organized around the main sulci. Such focal functional pulvinar projection fields were found in the lateral sulcus, the superior temporal sulcus, the intraparietal sulcus, the prefrontal cortex, the orbito-frontal cortex, the central sulcus and the anterior cingulate cortex. Although there are substantial inter-individual differences, these focal projection fields are consistently observed across hemispheres and multiple subjects. This calls for a reappraisal of the organization of pulvino-cortical functional connectivity loops.

## Material and methods

### Subjects

10 rhesus monkeys (Macaca mulatta) participated in the study (6 females, 4 males). Animal care procedures met all Belgian and European guidelines and were approved by the KU Leuven Medical School.

### Experimental setup

During the scanning sessions, monkeys sat in a sphinx position in a plastic monkey chair (Vanduffel et al., 2001) facing a translucent screen. Visual stimuli were retro-projected onto this translucent screen. Eye position (X, Y, right eye) was monitored thanks to a pupil– corneal reflection tracking system (RK-726PCI Iscan) at 120 Hz. During the resting state acquisitions, animals were required to maintain fixation into a 2×3 visual degrees tolerance window around a small red cross and were rewarded with apple juice for it.

### Scanning procedures

In this study, in-vivo MRI scans were performed with a 3T MR Siemens Trio scanner and PrismaFit in Leuven, Belgium.

#### Anatomical MRI acquisitions

Accompanying T1-weighted anatomical images were obtained during different sessions using a magnetization-prepared rapid gradient echo (MP-RAGE) sequence (TR=2200 ms, TE=4.06 ms, voxel size=0.4 mm by 0.4 mm by 0.4 mm). During the anatomical scans, the animals were sedated using ketamine/xylazine (ketamine 10 mg/kg I.M. 1 xylazine 0.5 mg/kg I.M., maintenance dose of 0.01–0.05 mg ketamine per minute I.V.).

#### Functional MRI acquisitions

Functional MRI acquisitions were as follows. Before each scanning session, a contrast agent, composed of monocrystalline iron oxide nanoparticles, Molday ION™ or Feraheme, was injected into the animal’s saphenous or femoral vein to increase the signal to noise ratio (Leite et al., 2002; Vanduffel et al., 2001). We acquired gradient-echo planar images covering the whole brain (40 slices, 84-by-84 in-plane matrix, flip angle=75°,repetition time (TR):2.00 s or 1.4s depending on the monkey; echo time (TE): 17-19 ms; resolution: 1.25×1.25×1.25 mm voxels) with an eight-channel phased-array receive coil; and a saddle-shaped, radial transmit-only surface coil (MRI Coil Laboratory, Laboratory for Neuro- and Psychophysiology, Katholieke Universiteit Leuven, Leuven, Belgium, see Kolster et al., 2014). Specific parameters varied across monkeys (Table 1).

**Table 1:**
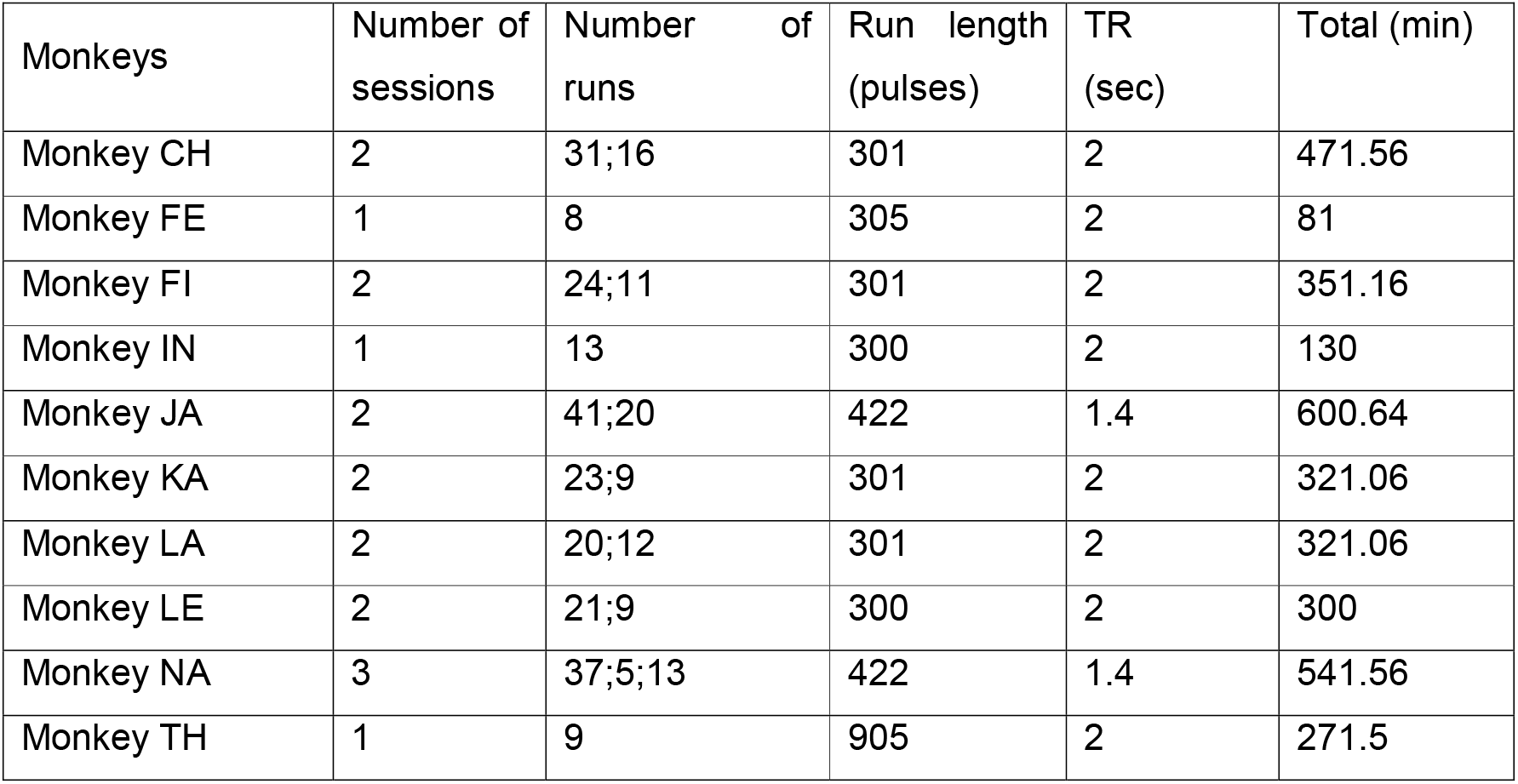
detailed acquisition parameters for each of the ten monkeys.

Functional volumes were corrected for head motion within and across sessions and slice time. They were linearly detrended, coregistered on the anatomical image and then normalized to the template space F99 with the resolution 1mm isotropic (http://sumsdb.wustl.edu/sums/macaquemore.do). A spatial smoothing was then applied with a 2-mm FWHM Gaussian Kernel.

### Data analysis

Runs were analyzed using AFNI (Cox, 1996) and FSL (FSL,RID:birnlex_2067; Jenkinson et al., 2012 http://fsl.fmrib.ox.ac.uk/fsl/fslwiki/).

The pulvinar of both hemispheres was drawn by hand for each monkey on their data registered and normalized to the F99 template (Figure 1). Individual functional voxels composing the pulvinar were defined as individual regions of interest. These ROIs were then used as seeds for a *seed-to-whole-brain analysis*. We performed this type of analysis for each ROI (voxel) and each run of each monkey. A Fisher’s r-to-z transformation was applied to the obtained correlation matrix. We then performed a t-test on the resulting maps of all the runs for each ROI. This statistical analysis resulted in a single correlation map (z-score) for each pulvinar ROI and each monkey, showing only the voxels that significantly correlated across all runs (p<0.001).

**Figure 1:**
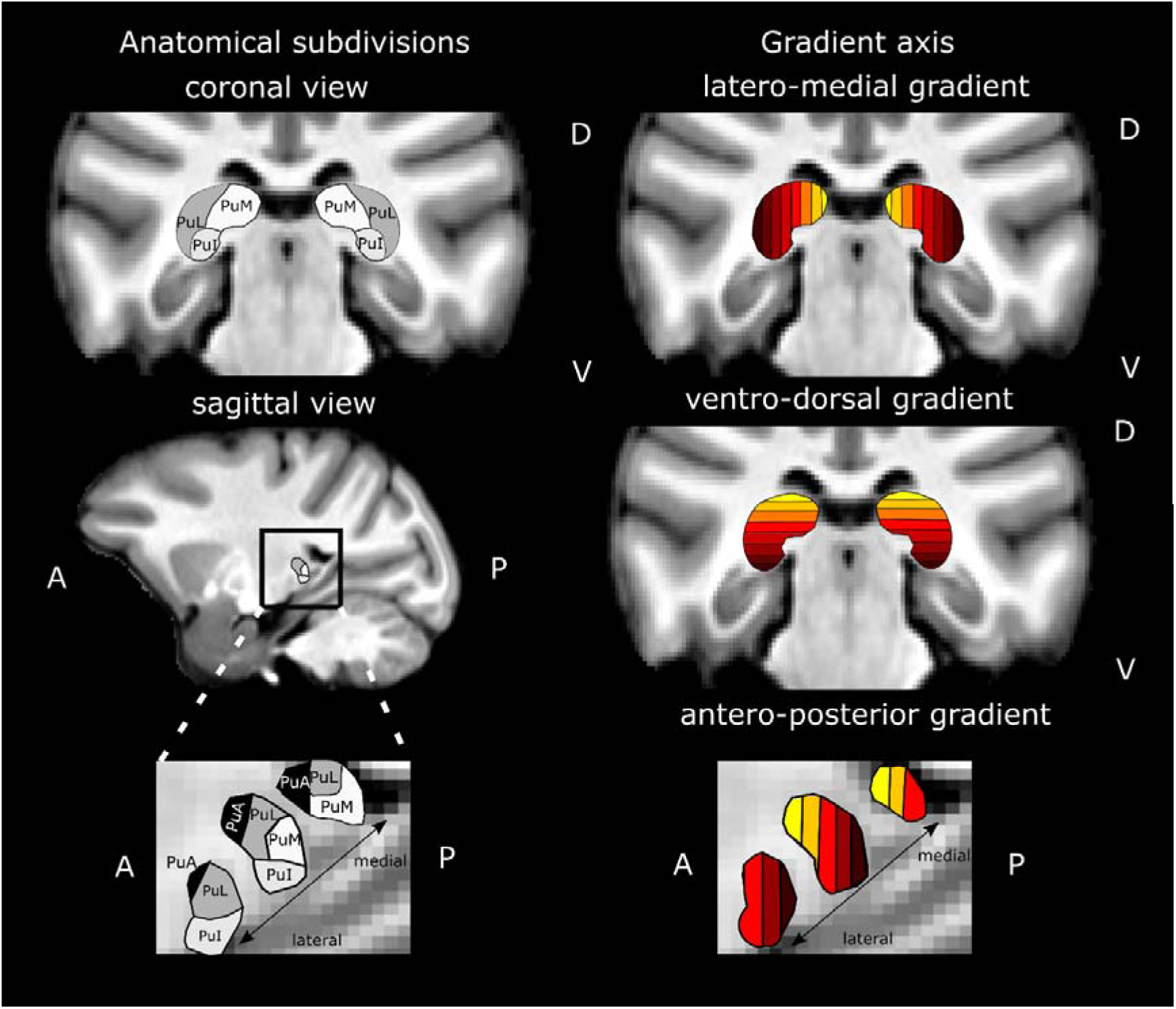
Anatomical pulvinar subdivisions and definition of latero-medial, ventro-dorsal and antero-posterior analysis gradients. The gradients are not based on anatomical subdivisions but can partially match with them depending on the chosen analysis gradient. For example, due to the shape of the pulvinar, PuI and PuM sub-regions will share the same color code along the ventro-dorsal gradient while PuM will be associated with lighter colors than PuL, along the latero-medial and antero-posterior gradients (See supplementary figure S1 for quantitative evaluations of this). PuL and PuA are respectively captured by the latero-medial gradient and antero-posterior gradient. PuI: inferior pulvinar; PuL: lateral pulvinar; PuM: medial pulvinar; PuA: anterior pulvinar.

All these correlation maps were compared to define for each voxel of the cortical map the pulvinar ROI with which the functional correlation value was highest (winner take all-within individual subjects). This “winner take all” analysis resulted in a single correlation map for each monkey where each cortical voxel was associated with the pulvinar ROI it correlated the most with (or no ROI if the statistical threshold in the z-score map was not reached). Thus, this map shows with high spatial resolution, the region of the pulvinar with which each cortical voxel correlates most. This is not an exclusive map, in the sense that it does not mean that cortical voxels do not also correlate with other pulvinar ROIs.

For visualization purposes, three color gradients have been created. The functional connectivity of pulvinar ROIs were grouped by slices defined along the latero-medial axis, the antero-posterior axis and the ventro-dorsal axis such that their color corresponded to a specific slice in the pulvinar along the specified axis (Figure 1). Each slice could thus include voxels from several sub parts of the pulvinar. The proportion of voxels belonging to each pulvinar sub parts, i.e. anterior, medial, lateral and inferior pulvinar, per slice are indicated in supplementary figure S1.

To observe the overall correlation across monkeys we computed a weighted average of the individual monkey correlation maps. To do so, we summed the maps across all monkeys and then divided the value of each voxel by the number of monkeys for which functional connectivity with the pulvinar was significant. In addition, we kept only voxels for which at least half of the monkeys presented a significant correlation.

To investigate the orientation of the pulvino-cortical connectivity at the local scale, i.e. at the scale of the sulcus, we attributed to each pulvinar slice or cortical location along a given sulcus a value describing its relative position along the ventral to dorsal (lower values for ventral and higher value for dorsal), the posterior to anterior, and the medial to lateral axes. We then performed linear regressions between the position within a given sulcus and the position of the most correlating pulvinar slices, across monkeys, for each hemisphere independently. This allowed us to investigate the spatial organization of the projections from the pulvinar to a given sulcus, addressing for example if the most dorsal slices of the pulvinar correlated most with the most dorsal slices of a given sulcus, or whether the most ventral slices of the pulvinar correlated most with the most ventral slices of this same sulcus. Supplementary table T1 summarizes the statistical outcome of these linear regressions for all selected sulci and both hemispheres.

## Results

The anatomical organization of the pulvinar in well-established sub-nuclei does not fully account for the functional organization of the pulvinar, nor for its functional connectivity with the cortex (reviewed in Froesel et al., 2021). Therefore, we investigated the topographical organization of pulvino-cortical functional connectivity. Ideally, one would like to describe pulvino-cortical functional connectivity at single-voxel resolution. However, this poses important visualization issues. In order to circumvent this, we decided for a more approximative approach. Specifically, we manually segmented the main pulvinar nuclei (figure 1, left panels), and we subdivided them in 2mm thick slices along the latero-medial (figure 1, top panel), ventro-dorsal (figure 1, middle panel) and antero-posterior axes (figure 1, bottom panel). We computed the functional connectivity of each of these slices with the rest of the brain and projected the winner-take-all voxels of this functional connectivity analysis on the corresponding semi-inflated brains. We used the same color code as in figure 1 (see methods; please note that only ipsilateral pulvino-cortical connectivity is considered). We first report a global organization of pulvino-cortical functional connectivity, which is observed at whole brain level and that can be precisely captured along the different axes described in figure 1. We then report local functional connectivity patterns that are observed in most individuals, though with some degree of inter-individual variability.

### Global pulvino-cortical functional connectivity gradients

The distinct functional connectivity patterns of the pulvinar with the cortex along the three aforementioned axes are presented in figure 2, for an individual subject (figure 2A) and averaged over all subjects (see methods, figure 2B). Individual subjects are present in Supplementary figure S2. The correlation maps represent the winner-take-all correlations between each voxel of the cortex and a unique pulvinar slice. This analysis is somewhat misleading as any given cortical voxel can be functionally connected to multiple pulvinar slices, but with varying functional connectivity strength. Lowering the criterion of maximal functional connectivity (winner-take-all approach) changed only marginally the reported observations (figure S3). This analysis leads to the description of complex functional connectivity gradients between the pulvinar and the cortex. It is important to note that due to the shape of the pulvinar, these gradients are not independent one from the other.

**Figure 2:**
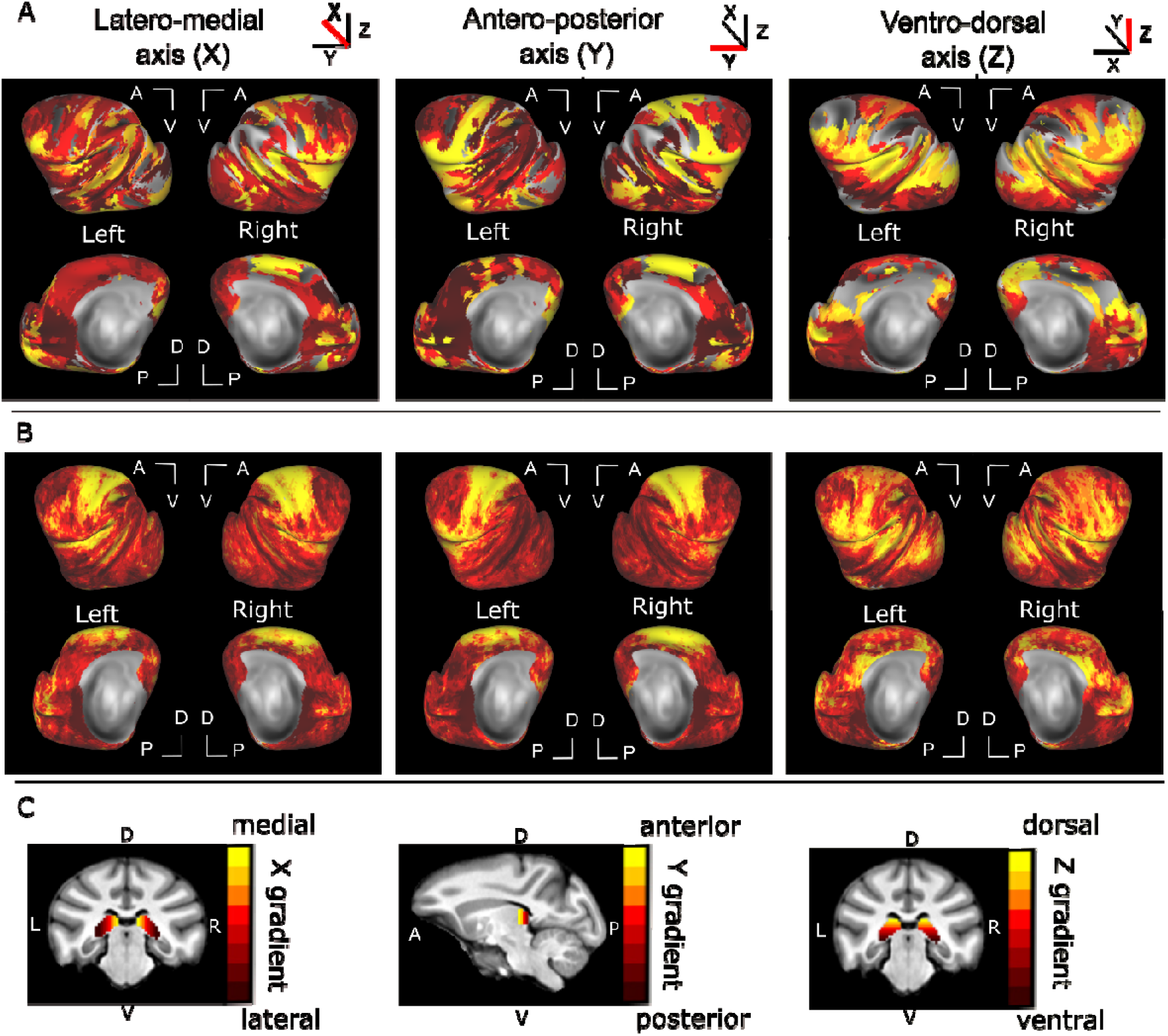
Global functional connectivity gradient of the pulvinar with the cortex. The pulvinar is subdivided into slices of 2mm thickness. These slices are used as seeds for a seed-to-whole brain functional connectivity analysis. A winner-take-all procedure is then applied to associate (color code) each cortical voxel with the pulvinar slice it maximally correlates with. Only ipsilateral correlations are presented. A) Winner-take-all functional connectivity map for a single subject. B) Mean winner-take-all functional connectivity maps across the 10 monkeys. C) Pulvinar slice seeds used in the functional connectivity analysis, for each of the analysis axes of interest. Details are provided in figure 1.

#### Latero-medial gradient

Both for a single subject (figure 2A, left panel, see also figure S2) and the group data (figure 2B, left panel), the sensorimotor cortex, the premotor cortex, the dorsal cingulate cortex and the insula are robustly correlated with the medial pulvinar. This is in line with the anatomical literature (Kaas & Lyon, 2007; Mufson & Mesulam, 1984; Padberg et al., 2009; Rosenberg et al., 2009; Wilke et al., 2018; Yeterian & Pandya, 1988). The medial pulvinar also shows reproducible functional connectivity with the inferior prefrontal cortex and the medio-posterior part of the superior temporal sulcus as well as a patchy connectivity with extrastriate cortex. In contrast, the lateral pulvinar has strongest functional connectivity with the anterior part of the superior temporal sulcus as well as with the dorsal lateral prefrontal cortex. These connectional features are better visible for individual brains and become less prominent in the group data, likely due to inter-individual differences. Note that this latero-medial parcellation of the pulvinar is partially overlapping with the anterior-posterior and ventral-dorsal parcellations, due to the specific 3D shape of the pulvinar.

#### Antero-posterior gradient

Similarly to the previous gradient, both on the single exemplar brain (figure 2A, middle panel, see also figure S2) and the group data (figure 2B, middle panel), the sensorimotor cortex, the premotor cortex, the dorsal cingulate cortex and the insula are functionally correlated with the anterior pulvinar (yellow) which is in line with the anatomical literature (Kaas & Lyon, 2007; Mufson & Mesulam, 1984; Padberg et al., 2009; Rosenberg et al., 2009; Wilke et al., 2018; Yeterian & Pandya, 1988). On the other hand, occipital areas and a large part of the posterior medial wall of the cortex are functionally more connected with the posterior pulvinar. Interestingly, the most anterior part of the brain, i.e. prefrontal and orbitofrontal cortex, shows stronger connectivity with the posterior and central parts of the pulvinar than with its anterior part, thus disrupting the antero-posterior gradient. Specifically, the prefrontal cortex correlates more with slices containing more medial pulvinar than anterior pulvinar, whether analyzing functional connectivity along the ventro-dorsal or the antero-posterior gradient (see Figure S1).

#### Ventro-dorsal gradient

In both single subject (figure 2A, right panel, see also figure S2,) and group data (figure 2B, right panel), strong functional connectivity can be observed between the dorsal part of the pulvinar and dorsal extrastriate cortex, the ventral part of the intraparietal sulcus, dorsal lateral sulcus, ventral sensorimotor cortex and ventral prefrontal cortex (yellow). More ventral pulvinar regions, on the other hand are functionally more connected with ventral occipital and temporal cortex. This ventro-dorsal functional gradient of the pulvino-cortical functional connectivity is also in agreement with the literature (Arcaro et al., 2015, 2018; Dominguez-Vargas et al., 2017; Shipp, 2003; Wilke et al., 2010).

A spatial correlation analysis between The “winner take all” spatial maps from the different subjects were significantly correlated across subjects (p < 0.001), indicating a consistency for the pulvino-cortical connectivity across subjects despite inter-individual differences (Spearman correlation coefficient (ρ): *antero-posterior gradient*: mean of both hemispheres = 0.46; *latero-medial gradient*: mean = 0.38; *ventro-dorsal gradient*: mean = 0.4). Overall, pulvino-cortical connectivity predominantly follows the ventro-dorsal and the antero-posterior gradient (except for the most anterior part of the cortex). It is interesting to note that the strongest correlation coefficients between subjects are reached by the antero-posterior gradient indicating a stronger inter-individual consistency along this axis (Friedman non-parametric test, X2(38) = 22.9, p < 0.001; post-hoc: antero-posterior axis vs latero-medial axis: p= 0.0001; antero-posterior axis vs ventro-dorsal axis: p= 0.0019; latero-medial axis vs ventro-dorsal axis: p=0.5938).

### Local connectivity gradients and multiple functional pulvino-cortical projection fields

While common global functional connectivity patterns can be identified between the pulvinar and the cortex characterized by both a ventro-dorsal and antero-posterior gradient, inter-individual differences do exist and some regions such as the prefrontal cortex escape these global functional connectivity patterns. This can be considered either noise, or a signature of specific local functional connectivity patterns between the pulvinar and the cortex. We addressed the latter hypothesis. In Figure 3, we focus on the functional connectivity of the pulvinar within restricted regions of the brain to investigate local connectivity gradients and to identify possible common functional connectivity patterns across different regions. Specifically, we focused on seven sulci and selected the functional connectivity axis of the pulvinar that resulted in statistically most robust topographic patterns (decision criterion: significant correlation in both hemispheres reflecting pulvino-cortical connectivity gradient along the same anatomical orientation, Supplementary Table 1).

**Figure 3:**
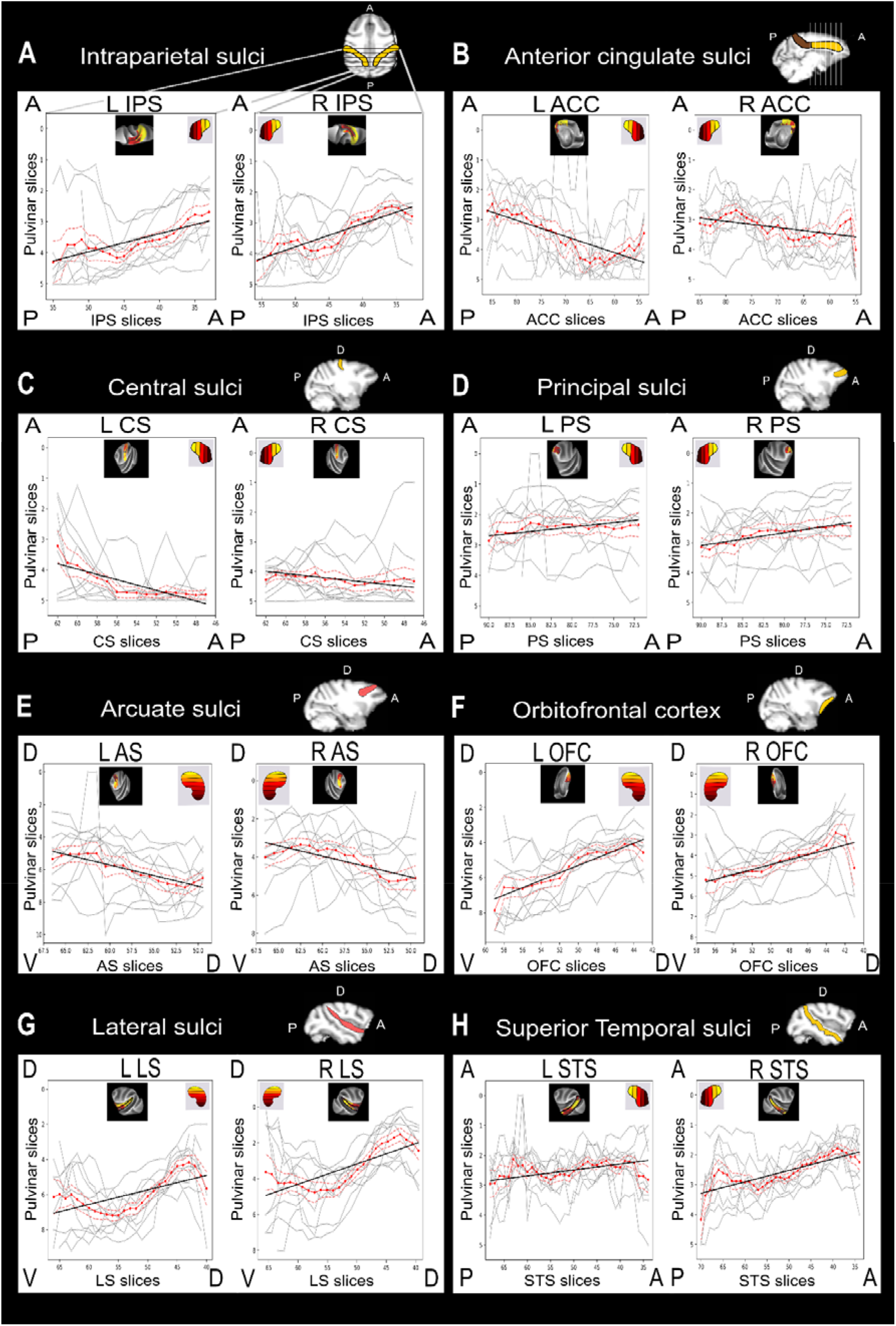
Local functional connectivity gradients of the pulvinar with specific brain regions. The pulvinar connectivity is retrieved for selected sulci and gyri: i.e., the intraparietal sulcus (A), anterior cingulate cortex (B), central sulcus (C), principal sulcus (D), arcuate sulcus (E), orbitofrontal cortex (F), lateral sulcus (G) and the superior temporal sulcus (H). Only ipsilateral correlations are presented. Sulci correlating with the pulvinar along the ventro-dorsal axis are colored in red and in yellow when the correlation follow the antero-posterior axis. These correlations are retrieved on the graph for each subject (grey) and the mean of the all the subjects (red) together with linear regressions (black). Note that only one anatomical axis is significant for both hemispheres for each sulcus.

#### Intraparietal sulcus

The intraparietal sulcus, which is mainly oriented along an antero-posterior axis, shows a functional connectivity pattern with the pulvinar, which follows an antero-posterior gradient, i.e. a gradient organized along the same axis as the sulcus (L IPS: p=19.9×10^-7^; r=0.33; R IPS: p=5.5×10^-14^; r=0.463, figure 3A, supplementary table S1).

#### Cingulate sulcus

The anterior cingulate cortex, which is also oriented along an antero-posterior axis, shows a pattern of functional connectivity with the pulvinar, again following an antero-posterior gradient, i.e. a gradient organized along the same axis as the sulcus (L ACC: p=2.4×10^-19^; r=0.46; R ACC: p=0.0002; r=0.285, figure 3B, supplementary table S1). However, contrary to the IPS, the ACC shows inverted connectivity with the pulvinar, whereby its most anterior part correlated more with the posterior sectors of the pulvinar. This indicates that the spatial organization of functional pulvinar projections to the cortex is not an artefact of distance from the nucleus. In comparison, the posterior cingulate cortex (PCC) does not present a clear orientation for its connectivity with the pulvinar (see supplementary table 1).

#### Central sulcus

The central sulcus, which is also oriented along an antero-posterior axis, shows a functional connectivity pattern with the pulvinar, again following an antero-posterior gradient, i.e. a gradient organized along the same axis as the sulcus (L CS: p=1,36×10^-12^; r=0.52; R CS: p=0.0004; r=0.27, figure 3C, supplementary table S1).

#### Prefrontal cortex

In the prefrontal cortex, the tip of the anterior principal sulcus was consistently connected with anterior pulvinar slices, and the pattern of functional connectivity gradually shifted towards more posterior pulvinar slices more posteriorly along this sulcus (L PS: p=0.04; r=0.36; R PS: p=0.001; r=0.23, figure 3D, supplementary table S1). The arcuate sulcus showed a reversed ventro-dorsal functional connectivity pattern with the pulvinar, with the pulvinar dorsal part correlating with the ventral most part of the arcuate sulcus (L AS: p= 2,336×10^-6^; r=0.46; R AS: p= 1.13×10^-3^; r=0.35, figure 3E, supplementary table S1). The orbitofrontal cortex demonstrated a ventro-dorsal connectivity pattern with the pulvinar (L OFC: p=7,06×10^-3^; r=0.51; R OFC: p=9,2×10^-13^; r=0.51, figure 3F, supplementary table S1). These diverse functional connectivity patterns in prefrontal cortex might explain why this region does not demonstrate a clear global functional connectivity pattern with the pulvinar.

#### Lateral sulcus

In the LS, the dorsal tip of the sulcus is mostly connected with dorsal pulvinar, and functional connectivity consistently shifted towards more ventral pulvinar slices as we moved anteriorly along the sulcus. This linear trend was disrupted when reaching the dorsal insula, anteriorly. This region expresses strong functional connectivity with the dorsal and medial pulvinar slices. A linear regression was performed to investigate the orientation of the pulvinar functional connectivity along these sulci. For both the left and right LS, the functional connectivity significantly followed a ventro-dorsal gradient (L LS: p=6.8×10^-12^; r=0.4; R LS: p=8.87×10^-14^; r=0.44, figure 3G, supplementary table S1).

#### Superior temporal sulcus

In the STS, the dorsal part of the sulcus is functionally connected most strongly with the dorsal pulvinar, and functional connectivity patterns consistently shifted to more ventral pulvinar slices as we moved anteriorly along the sulcus. This linear trend was disrupted when reaching the ventral part STS that shows a preferential connectivity with the dorsal pulvinar. This functional connectivity is in line with the literature showing functional connectivity between the most anterior AM face patch and the dorsal pulvinar (Schwiedrzik et al., 2015). Additionally, the medial pulvinar is functionally connected with the dorsal part of the STS and the lateral pulvinar with the ventral part of the STS. Thus, a dual functional connectivity gradient can be identified in this sulcus. As the STS shows this dual gradient along the ventro-dorsal axis (see supplementary figure S4), the linear regression along this axis was not significant (L STS: p=0.23; r=0.06; R STS: p=0.11; r=0.01), though the linear regression reached significance along the antero-posterior axis (L STS: p= 0,007; r=0.02; R STS: p= 1,06×10^-15^; r=0.4, figure 3H, supplementary table S1).

Overall, this suggests that, beyond the global gradients described in the previous sections, multiple pulvinar projection fields can be identified at cortical level. These projection fields, though systematic, are subject to substantial inter-individual variability and thus disappear on group average analyses. This observation, however, is expected to have profound implications on our understanding of the functional interactions between the pulvinar and the cortex.

## Discussion

Our results provide fine-grained functional connectivity patterns of the pulvinar with the cortex at several spatial scales. First, we show a global topographical connectivity pattern that can be captured along ventro-dorsal and antero-posterior pulvinar gradients and to a lesser extent along a medio-lateral gradient. We additionally describe more refined local functional connectivity patterns that capture multiple cortical pulvinar projection fields.

The goal of this work was to characterize more precisely the functional connectivity of the pulvinar with the cortex, in order to anatomically constrain its role in a variety of cognitive functions. In monkey fMRI studies, the cohort size is often a strong limitation (Milham et al., 2020, 2021). Here, we characterized pulvino-cortical functional connectivity patterns at rest, relying on ten awake fixating macaques. We confirm previous observations that the pulvinar has widespread connections with the cortex supporting its involvement in multiple cognitive functions (Barron et al., 2015; Froesel et al., 2021). Additionally, we show that the functional connectivity patterns we revealed resemble previously described anatomical connectivity thus describing a connectivity following a mix of the antero-posterior and ventro-dorsal axis (figures 4A and 4B present the functional connectivity as predicted by anatomical studies). These findings are in line with the neuronal recording and microstimulation literature (Kagan et al., 2020). We confirm the functional connectivity of the medio-dorsal pulvinar with attentional fronto-parietal networks (Fiebelkorn et al., 2019; Gutierrez et al., 2000; Kastner et al., 2020; Romanski et al., 1997). Indeed, the frontal eye field (FEF) and the lateral intraparietal area (LIP) represented in orange-yellow in both ventro-dorsal and latero-medial maps confirm their strongest functional correlations with the dorso-medial part of the pulvinar. The highly robust connectivity between the anterior part of the pulvinar and the sensorimotor and premotor cortex is also in agreement with previous studies (Kaas & Lyon, 2007; Mufson & Mesulam, 1984; Padberg et al., 2009; Rosenberg et al., 2009; Wilke et al., 2018; Yeterian & Pandya, 1988). The disruption of the anterior-posterior connectivity gradient in the prefrontal cortex is therefore also predicted by the literature, the anterior pulvinar being more connected with the motor and premotor regions whereas the medial pulvinar is described as interacting with the more anterior region of the brain such as the FEF, the DLPFC or the OFC (Froesel et al., 2021). Overall, the reported functional connectivity in the present work thus reliably reproduces observations collected with a diversity of methods and sets the stage for the analysis of pulvino-cortical functional connectivity at a more resolved spatial granularity.

**Figure 4:**
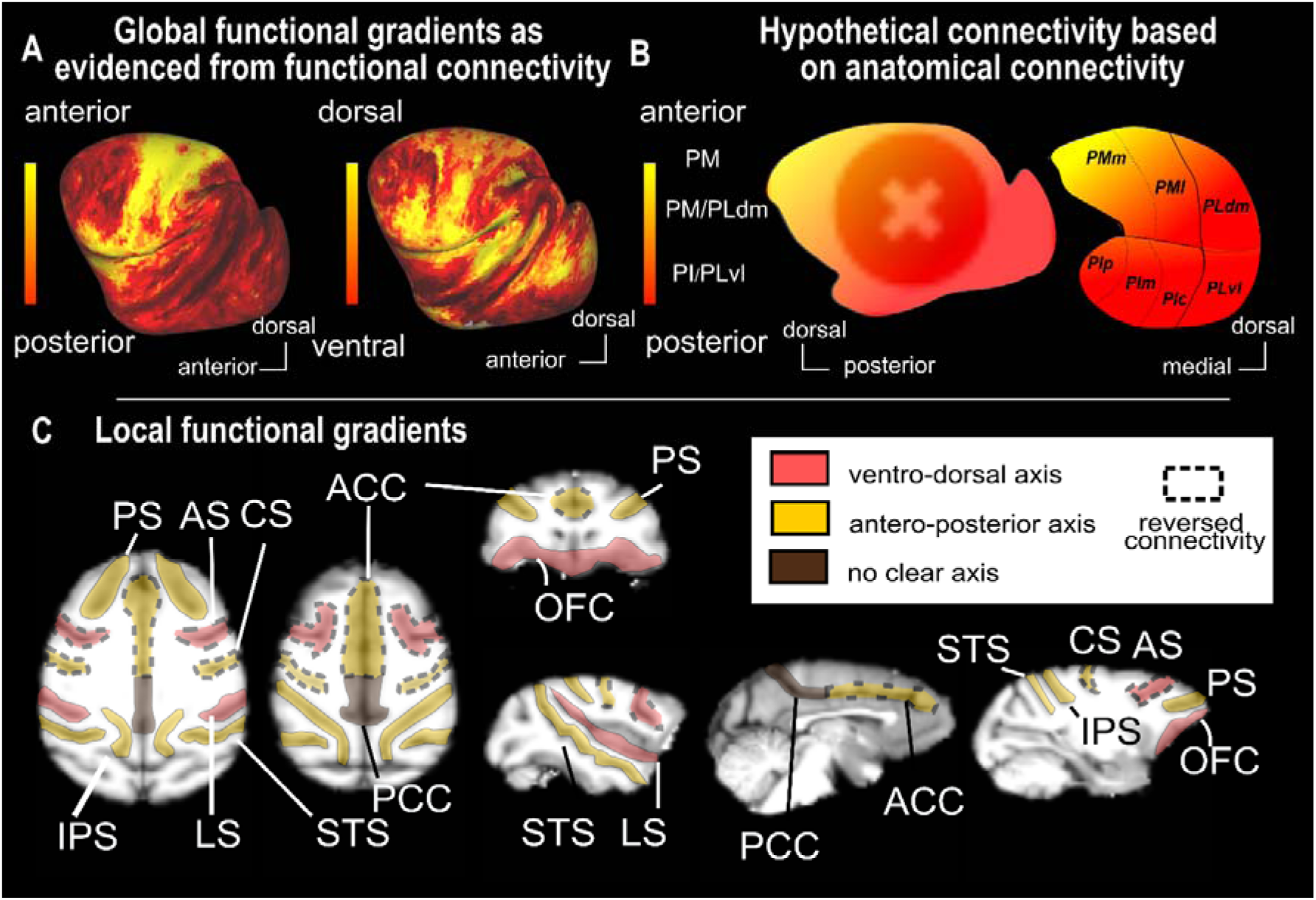
Summary of the global and local pulvino-cortical connectivity orientation. **A:** Results of the seed-to-whole brain functional connectivity analysis for the antero-posterior gradient (left) and the ventro-dorsal gradient (right), from figure 2. **B.** Attended mixed orientation of the functional connectivity based on the anatomical literature (based on the review Froesel et al., 2021). **C.** Summary of the local pulvino-cortical connectivity orientation. Dashed lines represented reversed connectivity pattern with the anatomical axis (ACC, AS and CS). PS: principal sulcus; ACC: Anterior cingulate sulcus; PCC: Posterior cingulate sulcus; OFC: orbitofrontal cortex; CS: central sulcus; AS: Arcuate sulcus; LS: lateral sulcus; STS: Superior temporal sulcus; IPS: intraparietal sulcus.

A major result of this work is that beyond the global connectivity gradients described above, multiple local pulvinar projections fields can be identified in multiple cortical regions. The organization of these pulvinar projections fields is reproduced across animals and hemispheres. Such local topographical organization of pulvinar connectivity has been occasionally reported in specific cortical regions such as the superior temporal sulcus or MT (Grimaldi et al., 2016; Mundinano et al., 2019), but to the best of our knowledge, never in such detail as here. We propose that this is a fundamental principle of pulvino-cortical connectivity and that a better description of these connectional principles will be crucial to fully understand the function of pulvino-cortical connections, as well as the role of the pulvinar in cognition. This organization, although preserved across individuals, is susceptible to some degree of inter-individual variability. We propose that this inter-individual variability that mostly takes place at the extremes of the sulci reflects morphological differences in the shape of the pulvinar and the sulci, which gets altered by the normalization onto the common reference brain. Supporting this interpretation, the segmentation of the pulvinar onto the individual brains improved the identification of the local pulvinar projection fields.

The local pulvinar projection fields are summarized in figure 4C. Using a color code, we indicated the dominant orientation of the functional connectivity patterns between the pulvinar and the different sulci. The orientation of the functional connectivity gradients is dominated by ventro-dorsal and antero-posterior pulvinar projections, often matching the dominant orientation of the sulci. It is interesting to highlight that the sulci from the posterior part of the brain that tend to follow the antero-posterior axis, matching the global gradient, while the sulci from the prefrontal region show more diverse preferred orientations of the connectivity. Please also note that while the color reflects dominant functional connectivity orientation, it does not indicate the polarity of the projections. For example, the anterior cingulate cortex has a reversed antero-posterior pulvinar mapping relative to the other sulci dominated by antero-posterior pulvinar projections. This reversed connectivity pattern is also found in the central sulcus and the arcuate sulcus, possibly suggesting a very specific and more refined organization of the connectivity between the pulvinar and the anterior part of the cortex ad compared to the posterior part of the brain. The dual functional connectivity gradient double gradient observed within the STS, is yet another expression of the complex organization of the pulvino-cortical functional connectivity. While this sulcus follows a global antero-posterior gradient of functional connectivity with the pulvinar two ventro-dorsal gradients are observed thus determining to pulvinar projections field along the STS. We hypothesize that this rich connectivity allows a very refined interaction between the subcortical nucleus and the superior temporal sulcus. Fine multimodal studies combining high resolution fiber tracking (Tounekti et al., 2018) and voxel-based functional connectivity analysis are expected to clarify the origin of these multiple cortical pulvinar projection fields.

The description of these functional gradients characterizing the local cortical pulvinar projection fields does not exclude the possibility that any given cortical region can receive input from multiple pulvinar sectors. For example, a correlation of the STS and others cortical regions can be described with both ventral and dorsal pulvinar (data not shown), confirming the microstimulation results recently reported by Kagan and colleagues (Kagan et al., 2020). These projections from multiple pulvinar nuclei in specific brain regions has been a core argument in support of the role of the pulvinar as a modulator (Benarroch, 2015; Saalmann & Kastner, 2011). The pulvinar can thus be a part of several networks and can be recruited differentially as a function of sensory modality or the ongoing cognitive demand (Froesel et al., 2022), thus supporting cognitive flexibility. As a result, another important aspect to keep in mind is that the winner-take-all maps we report here from monkeys at rest could be different during active tasks involving either sensory stimulation or the production of complex behaviors. We hypothesize that at rest, the winner-take-all maps will follow more closely the anatomical structural connectivity (Honey et al., 2007, 2009), while in contrast, during systemic localizer studies (Russ et al., 2021) or active behavior, the correlations would be more pronounced in specific functional networks, the functional connectivity with other subregions being possibly attenuated. Last, when interpreting the current observations, technical aspects need to be kept in mind in addition to the interpretation of the winner-take-all approach and the possibility of overlapping connectivity fields from different pulvinar subregions. Some of these aspects pertain to the spatial resolution of the data and associated possible partial volume effects, possibly obscuring a finer grained anatomical organization. Other aspects pertain to the very measure of functional connectivity from resting state data used here, that lacks directionality information and that might occasionally describe second order rather than first order connectivity. Fine-grained multimodal studies combining rs-fMRI, DTI and anatomical tracing will be needed to resolve these issues.

## Conclusion

Two scales of functional connectivity between the pulvinar and the cortex are described. A global topographical functional connectivity pattern that can mainly be captured along the antero-posterior and ventro-dorsal axes, and multiple local topographically organized pulvinar projections fields in multiple cortical regions such as the lateral sulcus, the superior temporal sulcus, anterior cingulate cortex and the intraparietal sulcus. In spite of the inter-individual differences, the local functional gradients are consistently described across individuals and in both hemispheres and follow the anatomical orientation of the sulci. We propose that these multiple pulvinar projection fields correspond to a fundamental principle of pulvino-cortical connectivity and that a better understanding of this organization will help clarifying the function of pulvino-cortical connectivity and functions.

## Author contribution

Conceptualization: S.B.H., M.F; Data Curation: W.V., M.F., Q.Z.; Formal Analysis: M.F., Q.G. and S.B.H.; Funding Acquisition: S.B.H.; Investigation: M.F., and S.B.H.; Methodology: M.F., Q.G., C.S., and S.B.H.; Resources: Q. Z. and W.V.; Supervision: S.B.H.; Validation: S.B.H.; Visualization: M.F., C.S., Q.G.; Writing-Original draft: M.F. and S.B.H.; Writing-review & editing: M.F., C.S., Q.G., Q. Z., W.V., and S.B.H.

## Ethical statement

Animal care and experimental procedures were performed in accordance with the National Institute of Health’s Guide for the Care and Use of Laboratory Animal, the European legislation (Directive 2010/63/EU) and were approved by the Animal Ethics Committee of the KU Leuven. Weatherall reports were used as reference for animal housing and handling. All animals were group-housed in cages sized 16-32 m^3^, which encourages social interactions and locomotor behavior. The environment was enriched by foraging devices and toys. The animals were fed daily with standard primate chow supplemented with fruits, vegetables, bread, peanuts, cashew nuts, raisins and dry apricots. The animals were exposed to natural light and additional artificial light for 12 h every day. On training and experimental days, the animals were allowed unlimited access to fluid through their performance during the experiments. Using operant conditioning techniques with positive reinforcers, the animals received fluid rewards for every correctly performed trial. During non-working days, they received water in their living quarters. Throughout the study, the animals’ psychological and veterinary welfare was monitored daily by the veterinarians, the animal facility staff and the lab’s scientists, all specialized in working with non-human primates. The animals were healthy at the conclusion of our study. M1 and M2 are currently still employed in other studies.

## Acknowledgments

S.B.H. was funded by the French National Research Agency (ANR) ANR-16-CE37-0009-01 grant and the LABEX CORTEX funding (ANR-11-LABX-0042) from the Université de Lyon, within the program Investissements d’Avenir (ANR-11-IDEX-0007) operated by the French National Research Agency (ANR). This work received funding from KU Leuven C14/21/111, IDN/20/016, C3/21/027; Fonds Wetenschappelijk Onderzoek-Vlaanderen (FWO-Flanders) G0E0520N, G0C1920N. We thank also thank Thomas Perret, Johan Pacquit and Marco Bimbi for help with the hardware computational resources.

## Supplementary data

**Supplemental Table 1:**
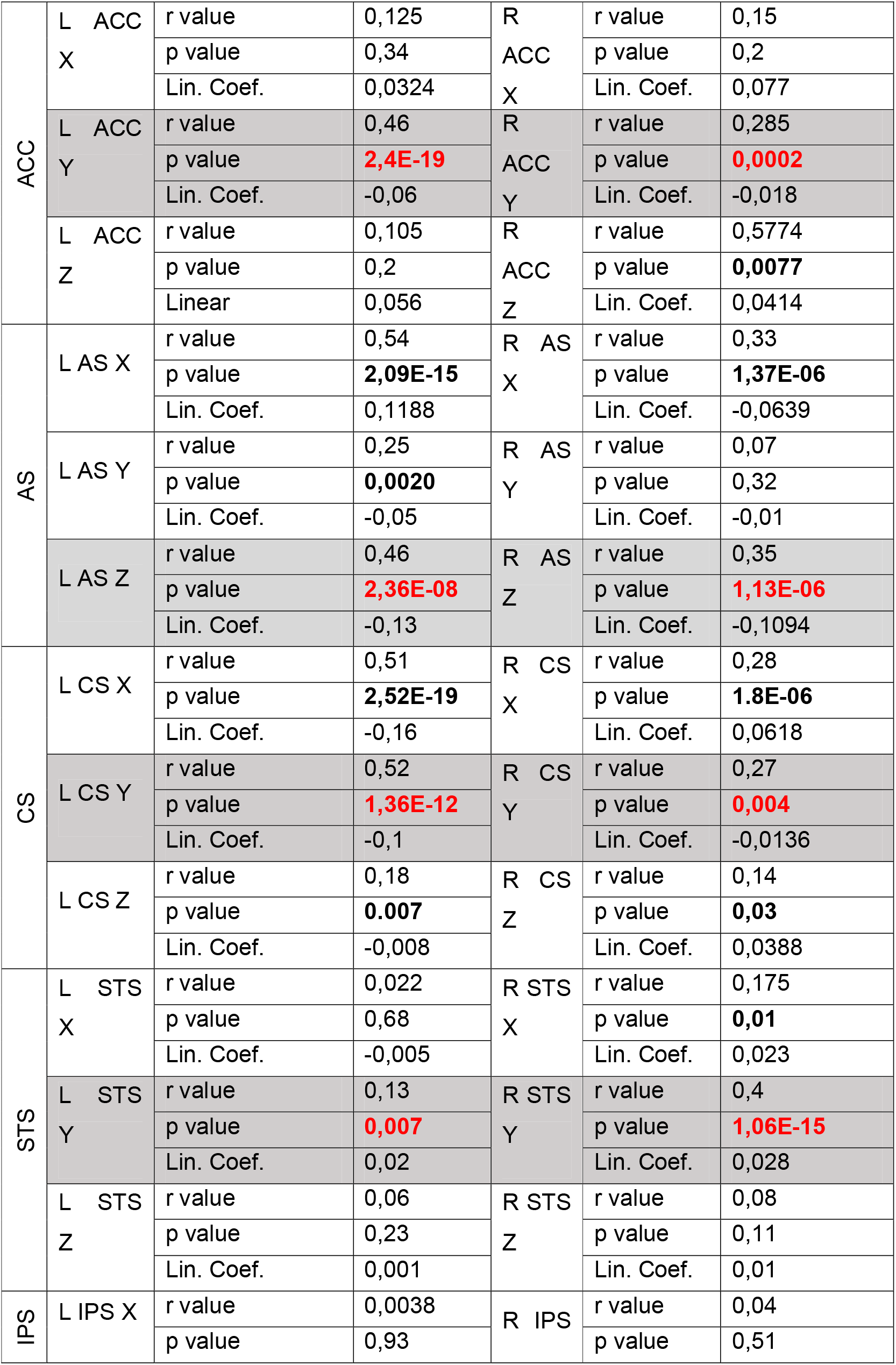
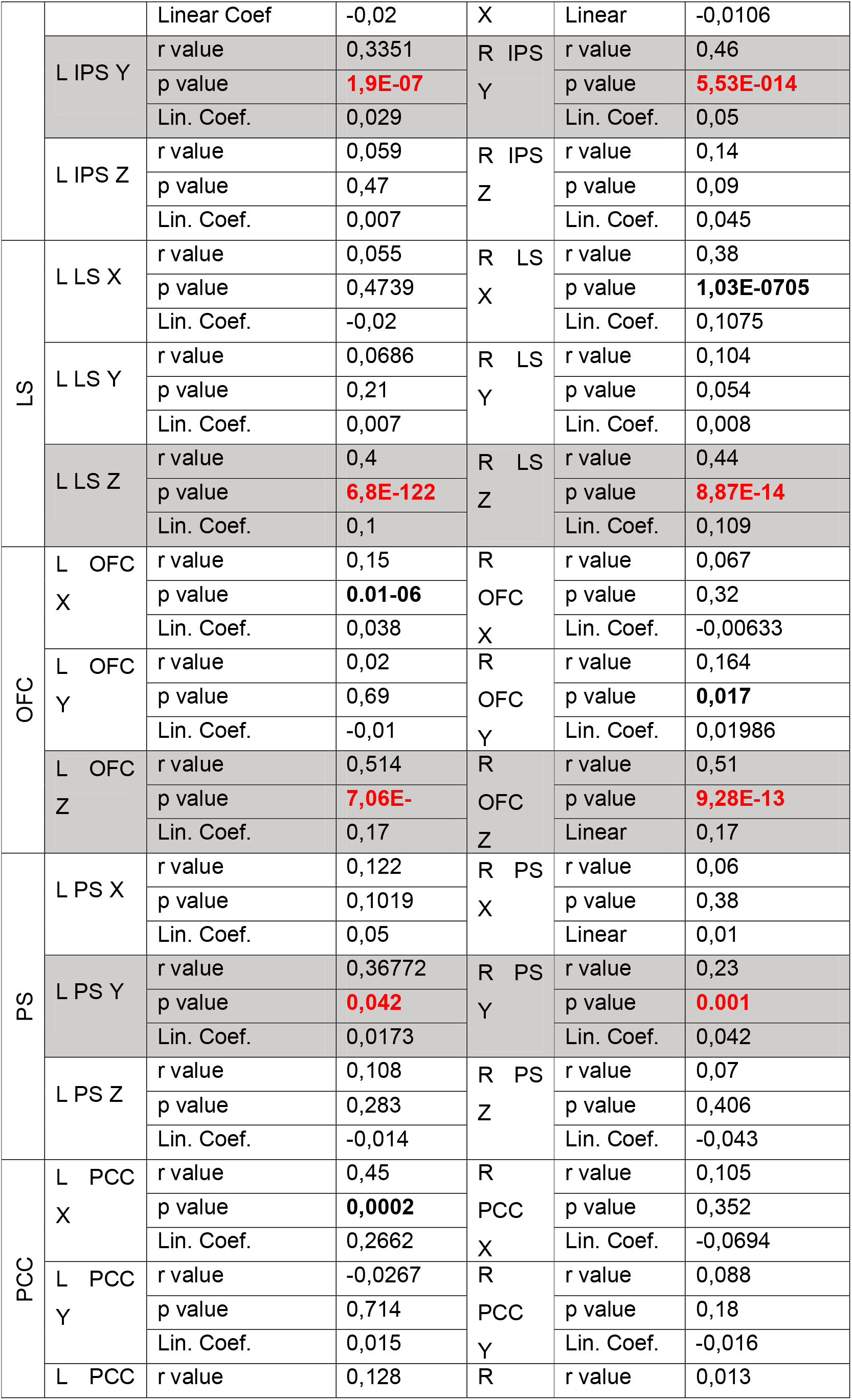
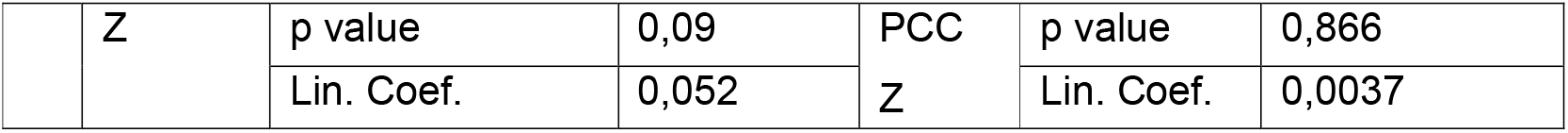
Detailed statistics of the linear regression analysis. Significative correlations are indicated in bold. Anatomical orientation presenting a significant correlation in both hemispheres with the same orientation for any given sulcus is indicated in bold red and highlighted against a grey background. X: latero-medial axis. Y: antero-posterior axis. Z: ventro-dorsal axis.

**Figure S1:**
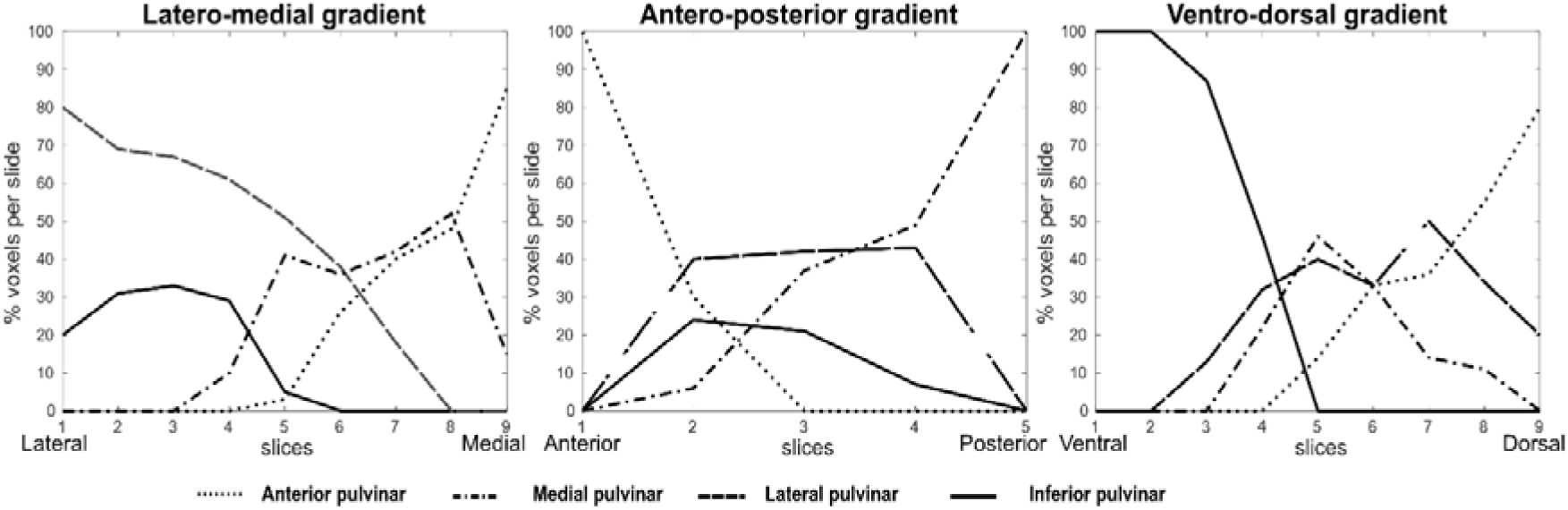
Proportion of voxels belonging to each pulvinar sub parts, i.e. anterior, medial, lateral and inferior pulvinar, per slice.

**Figure S2:**
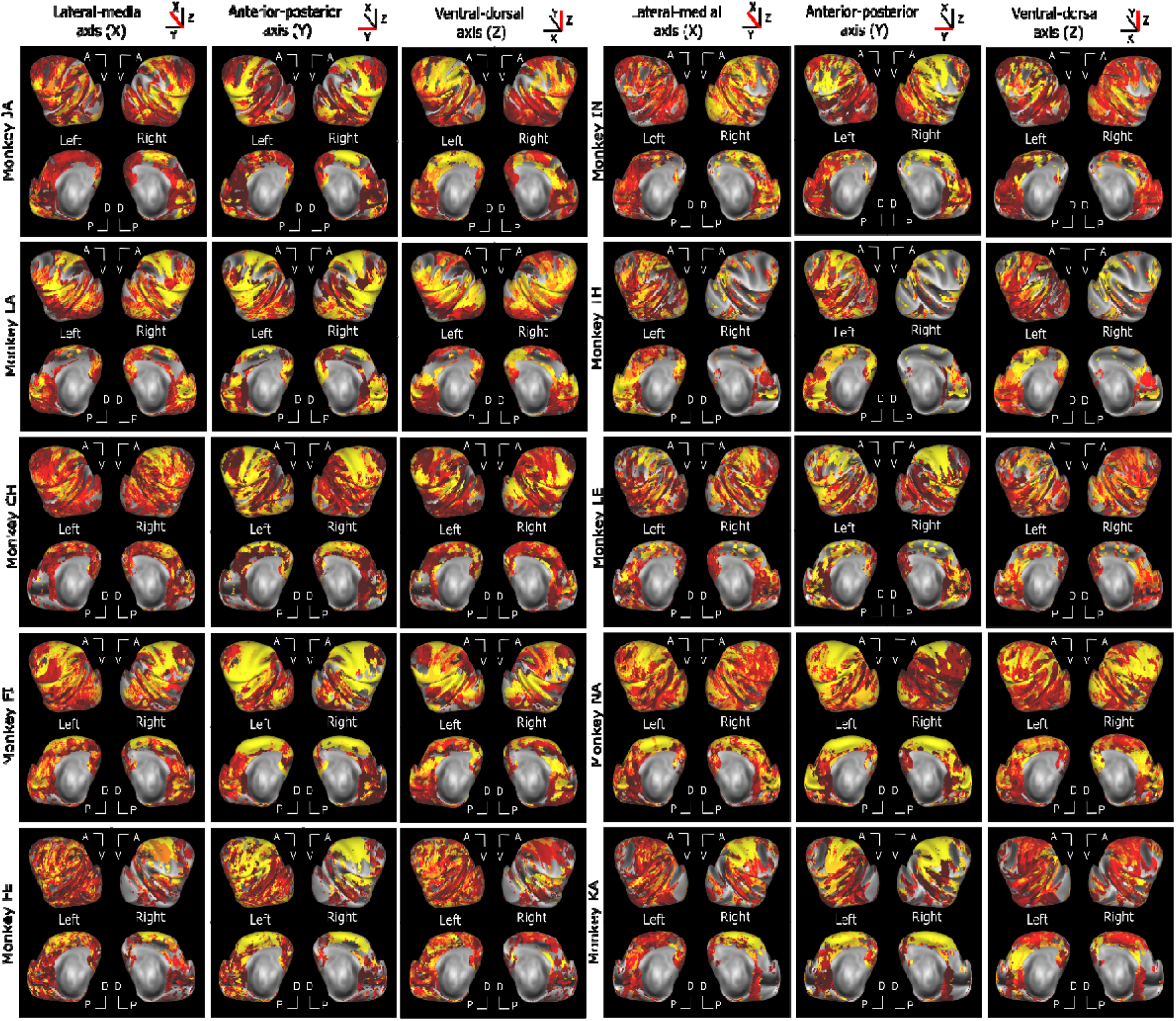
Winner take all functional connectivity maps for each individual monkey. All else as in figure 2. Only ipsilateral correlations are presented.

**Figure S3:**
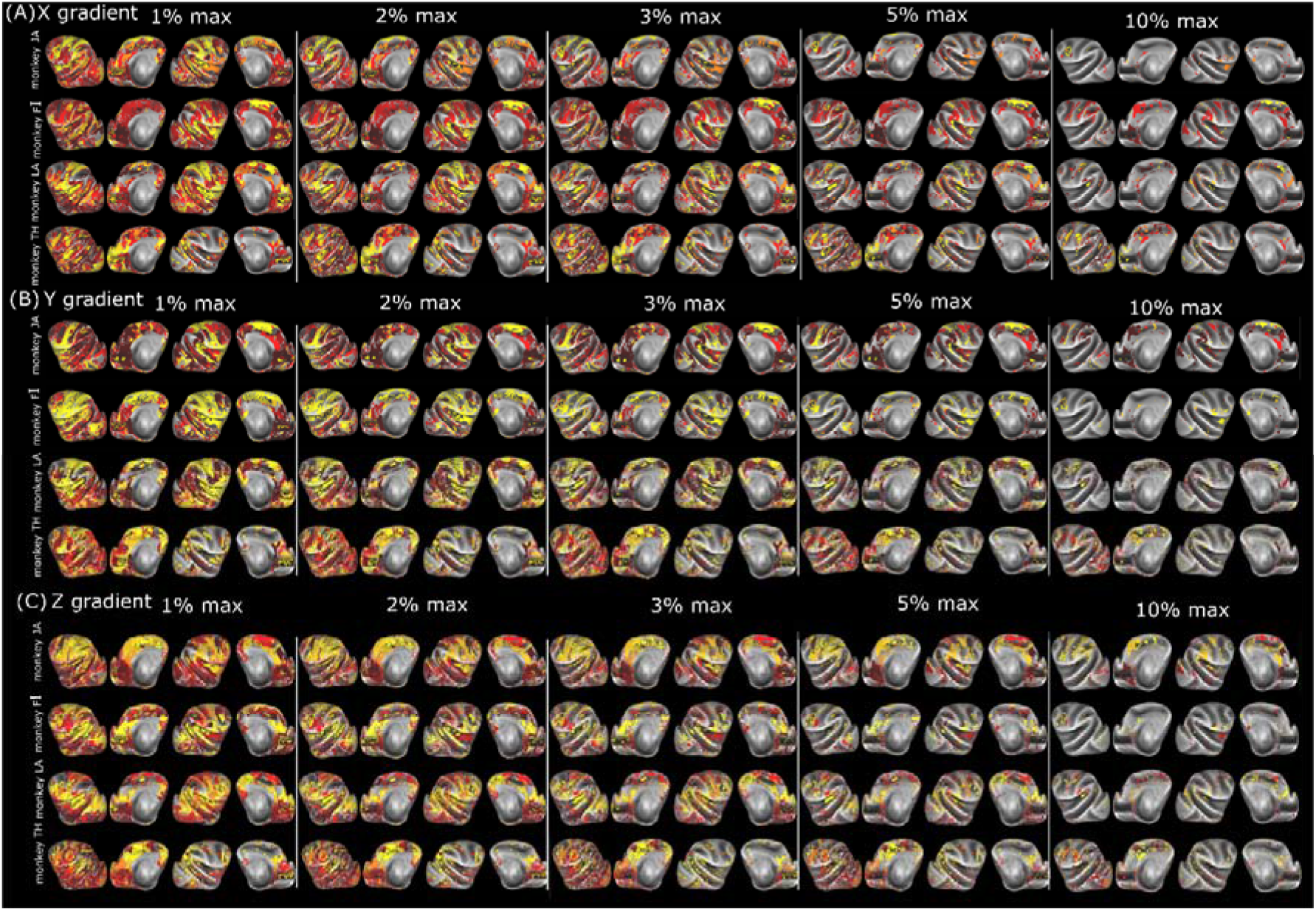
Winner take all functional connectivity maps for individual monkeys as a function of percent of correlation strength distance to the second best correlating voxel. The maps are the winner take all functional connectivity maps displaying only the correlation that are 1, 2,3, 5 or 10% higher than the second best correlation. Four exemplar monkeys. Only ipsilateral correlations are presented.

**Figure S4:**
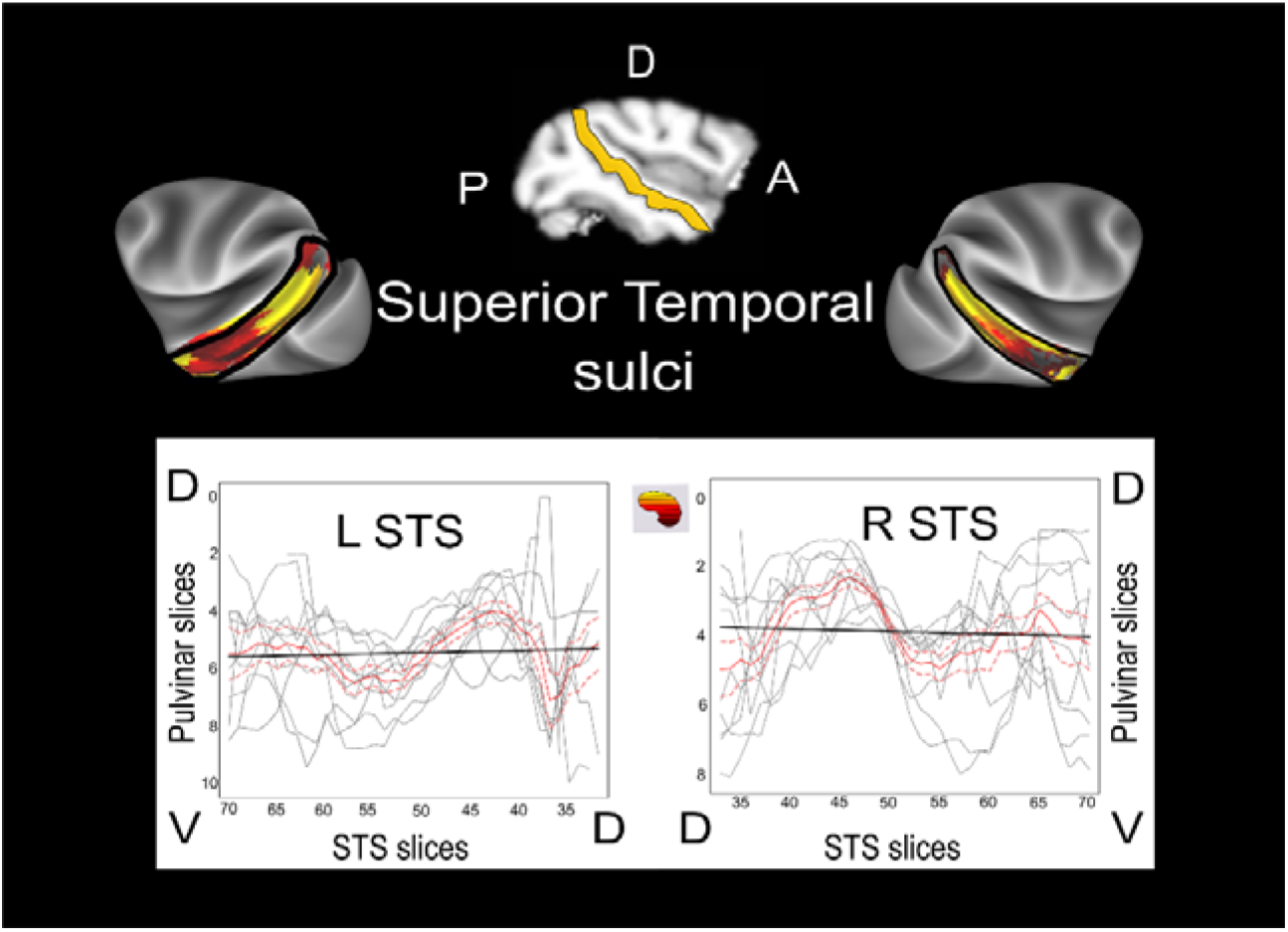
Local functional connectivity gradients of the pulvinar with the superior temporal sulcus. Only ipsilateral correlations between the STS and the pulvinar are presented for the STS along the ventro-dorsal axis. Individual subject (grey) and mean across all the subjects (red) correlations are presented as well as the group linear regression (black).

